# CitrusKB: A Comprehensive Knowledge Base for Transcriptome and Interactome of *Citrus* spp. Infected by *Xanthomonas citri* subsp. *citri* at Different Infection Stages

**DOI:** 10.1101/2020.03.19.997999

**Authors:** Adriano Ferrasa, Mayara M. Murata, Teresa D. C. G. Cofre, Juliana S. Cavallini, Gustavo Peron, Maria H. M. Julião, José Belasque, Henrique Ferreira, Maria Inês T. Ferro, Rui P. Leite, Helen A. Penha, Flávia M. S. Carvalho, Alessandro M. Varani, Roberto H. Herai, Jesus A. Ferro

**Affiliations:** Departamento de Informática, Universidade Estadual de Ponta Grossa (UEPG), Ponta Grossa, Paraná, Brazil; Departamento de Tecnologia, Universidade Estadual Paulista (UNESP), Jaboticabal, São Paulo, Brazil; Graduate Program in Health Sciences, School of Medicine, Pontifícia Universidade Católica do Paraná (PUCPR), Curitiba, Paraná, Brazil; Departamento de Fitopatologia e Nematologia, Escola Superior de Agricultura “Luiz de Queiroz”, Universidade de São Paulo (USP), Piracicaba, São Paulo, Brazil; Departamento de Biologia Geral e Aplicada, Instituto de Biociências, Universidade Estadual Paulista (UNESP), Rio Claro, São Paulo, Brazil; Instituto Agronômico do Paraná (IAPAR), Londrina, Paraná, Brazil

**Keywords:** *Citrus* spp., plant-pathogen interaction, citrus canker, transcriptome sequencing, differential gene expression, web-based interface

## Abstract

Citrus canker type A is a serious disease caused by *Xanthomonas citri* subsp. *citri* (*X. citri*), which is responsible for severe losses to growers and to the citrus industry worldwide. To date, no canker-resistant citrus genotypes are available, and there is limited information regarding the molecular and genetic mechanisms involved in the early stages of the citrus canker development. Here, we present the knowledge base for transcriptome of *in vivo* citrus interactome, the CitrusKB. This is the first *in vivo* interactome database for different citrus cultivars, and it was produced to provide a valuable source of information on citrus and their interaction with the citrus canker bacterium *X. citri*. The database provides tools for a user-friendly web interface to search and analyze a large amount of information regarding eight citrus cultivars with distinct levels of susceptibility to the disease and their interaction, at different stages of infection, with the citrus canker bacterium *X. citri*. Currently, CitrusKB comprises a reference citrus genome and its transcriptome, expressed transcripts, pseudogenes and predicted genomic variations (SNPs and SSRs). The updating process will continue by incorporating annotations and analysis tools. We expect that CitrusKB may substantially contribute to the area of citrus genomics. CitrusKB is accessible at http://bioinfo.deinfo.uepg.br/citrus. Users can download all the generated raw sequences and generated datasets by this study from the CitrusKB website.

## Background

Citrus canker A, caused by the Gram-negative bacterium *Xanthomonas citri* subsp. *citri* (*X. citri*), is one of the main diseases affecting citrus trees and is a threat for orange production in several countries around the world [1–3]. The symptoms of citrus canker on susceptible trees include raised brownish circular lesions on leaves, stems and fruits [4]. Several disease management procedures have been applied in attempts to control citrus canker, including pruning of infected trees, pos-harvested treatment of fruits, decontamination of equipment and personnel and, within endemic regions of citrus canker, it is applied a spray of copper-containing chemicals to protect young leaves and fruit against bacterium infection [5, 6]. Furthermore, additional measures to control the disease involves planting windbreaks in the citrus orchards, control of the citrus leaf miner *Phyllocnistis citrella*, and production of healthy citrus nursery trees [6, 7].

Due to these challenging attempts for disease control, several efforts have been made to understand the mechanisms of plant-pathogen interaction and the disease development molecular basis, with the objective to establish more effective measures for control of citrus canker.

The molecular basis and the genetic mechanisms involved in the early stages of the citrus canker development may be revealed by studies on citrus species and cultivars with different levels of susceptibility to the disease. For instance, kumquats (*Fortunella* spp.) and ‘Mexican’ lime (*Citrus aurantifolia* (Christm.) Swingle) are considered resistant and susceptible to citrus canker, respectively. Furthermore, other citrus cultivars exhibit intermediate levels of susceptibility and resistance to the disease, such as the sweet oranges (*Citrus sinensis* L. Osbeck) ‘Bahia’, ‘Hamlin’, ‘Valencia’, and ‘Pera’, and the mandarins ‘Ponkan’ (*Citrus reticulata* Blanco) and Satsuma (*Citrus unshiu* Marcovitch) [7, 8] (Figure 1). Therefore, the study of the biological processes of the plant-pathogen interaction at molecular level in such diverse citrus genotypes may help to better understand citrus canker disease and may allow identifying potential targets for disease management. For instance, the high throughput RNA-Seq technology can be used to detect the induced or repressed genes in the host during the bacterial infection process. Then, the transcriptome can unveil the genetic and molecular mechanisms that confer the different levels of citrus susceptibility and resistance against the bacterial pathogen. However, current transcriptome-based studies did not include a large range of citrus cultivars exhibiting different levels of resistance and susceptibility to citrus canker. For instance, a previous study was carried out using microarray-based analysis in sweet oranges ‘Pera’ and ‘Cristal’ varieties (*C. sinensis*) and ‘Mexican’ lime (*C. aurantifolia*) and ‘Siciliano’ lemon (*Citrus limon* (L.) Burm. f.) [9]. In another investigation, based on RNA-Seq analysis, only two cultivars, the canker-resistant ‘Meiwa’ kumquat (*Fortunella crassifolia* Swingle), and the canker-susceptible ‘Newhall navel sweet orange’ (*C. sinensis*) were compared [10]. In addition, RNA-Seq technique was also used to develop a high confident reference of citrus transcriptome based on 12 *Citrus* species from all main phylogenetic groups [11].

**Figure 1.**
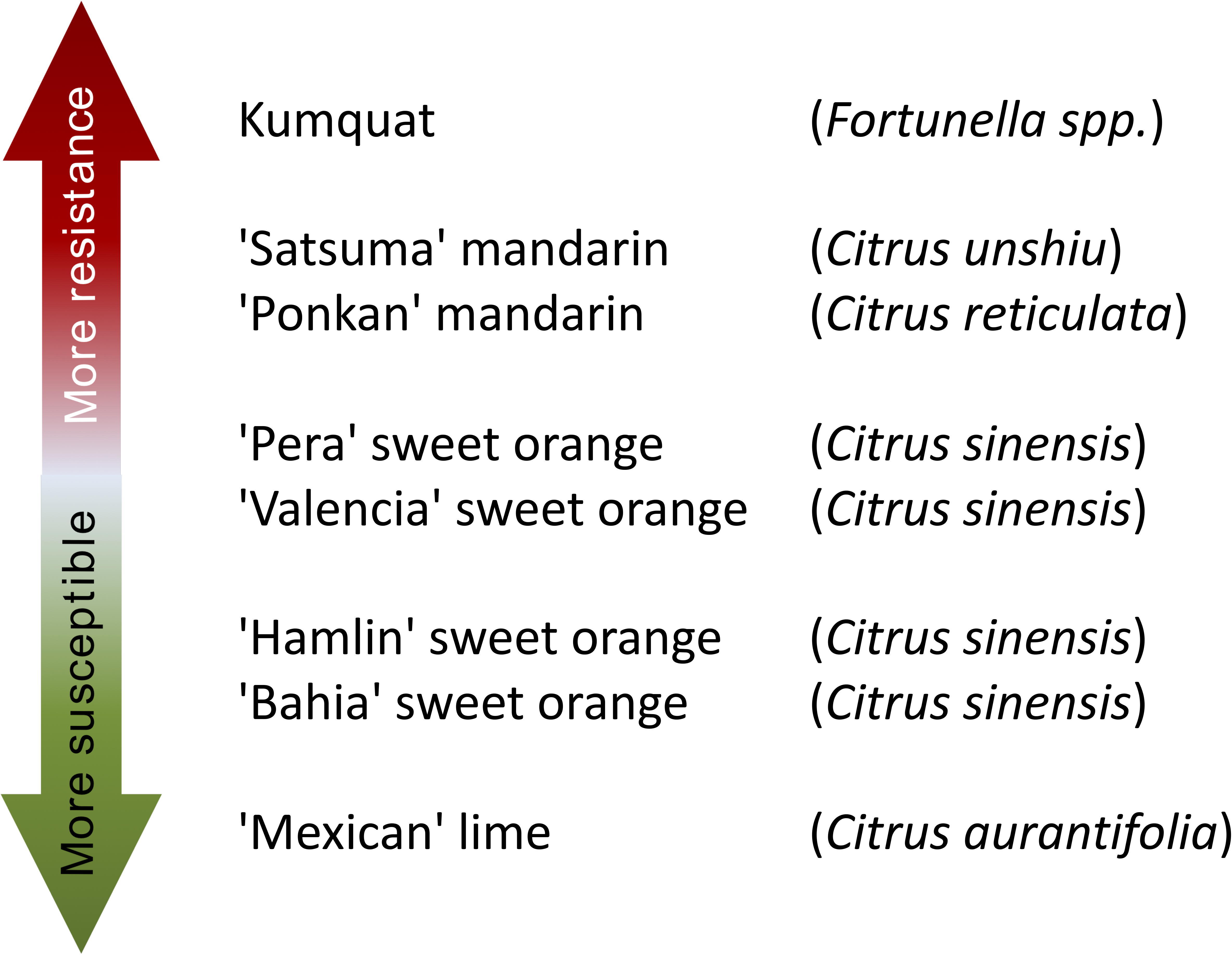
Citrus genotypes resistance and susceptibility scale to citrus canker A. ‘More resistant’ indicates that a citrus genotype is more resistant to citrus canker disease, as well as ‘More susceptible’ indicates that a citrus genotype is less resistant to citrus canker disease.

In this study, we aimed to determine early genetic expression changes in citrus plants under the development of the citrus canker disease using RNA-Seq approach to sequence the expressed mRNA of eight different citrus genotypes inoculated with *X. citri* at early stages of the infection by the bacterium *X. citri*. All sequenced data were analyzed to detect the differentially expressed genes between compared genotypes. Thus, we have developed a transcriptome knowledge base of distinct citrus cultivars, the CitrusKB, to facilitate further research on the biology of citrus. Despite other citrus databases, such as *Citrus sinensis* annotation project [12] and Citrus Genome Database (CGD, https://www.citrusgenomedb.org), CitrusKB is the only web tool that comes to integrate a wide range of citrus species and cultivars exhibiting different levels of resistance to citrus canker. The CitrusKB is a user-friendly web-based interface that provides tools to allow researchers to visualize, search, analyze, recover and browse information on citrus and *X. citri* interactome.

## Construction and Content

Currently, CitrusKB hosts a database to study the effects of initial stages of citrus canker disease including *in vivo* RNA-Seq data of eight citrus genotypes, which have been prepared during three stages, at 24, 48 and 72 hours post-inoculation (hpi) of *X. citri*. In addition, the website includes the sequences of other citrus reference genomes that were available in other databases.

### Inoculation of citrus genotypes with *X. citri*

The eight citrus genotypes exhibiting different levels of resistance to citrus canker included in the study were: ‘Kumquat’ (*Fortunella* spp.); Mandarins ‘Ponkan’ (*Citrus reticulata*) and ‘Satsuma’ (*Citrus unshiu*); sweet oranges (*Citrus sinensis*) ‘Bahia’, ‘Valencia’, ‘Pera’, and ‘Hamlin’; and ‘Mexican’ lime (*Citrus aurantifolia*). The trees were maintained in a greenhouse under controlled temperature (28°C) and photoperiod conditions (light/dark periods of 12 hours). The trees were pruned three weeks prior to inoculation in order to produce uniform young leaves, with 75% of leaf expansion (ideal condition for bacterial inoculation test in citrus [13]). The citrus trees were grafted on ‘Rangpur’ lime (*Citrus limonia* Osbeck). The *X. citri* strain 306 [14] was streaked from glycerol stock on Nutrient Agar (0.3% beef extract, 0.5% peptone, 1.5% agar) for 72 h at 28 °C. A single colony was transferred to a new Nutrient Agar plate and after 72 h at 28 °C, a bacterial sample was diluted into sterile bi-distilled water and the inoculum was prepared by dilution to 10^8^ cfu/ml (0.3 OD reading at 600 nm). The bacterial suspension was infiltrated into the abaxial surface of five leaves of each of the three plants in the entire leaf blade by using a 1 mL syringe without needle. Similarly, leaves of other three check plants were infiltrated with sterile distilled water (control).

### Citrus RNA extraction and sequencing

Treated and control plants were kept in a growth chamber under controlled temperature (28°C) and photoperiod conditions (light/dark periods of 12 hours). At 24, 48 and 72 hours after inoculation, three leaves inoculated with *X. citri* and showing canker symptoms (infected - I) and three leaves inoculated with sterile water and without canker symptoms (control -NI) were collected from each plant and immediately frozen in liquid nitrogen and further stored at −80°C.

For each time, frozen leaf tissue was ground with mortar and pestle using liquid nitrogen. Total RNA was extracted from macerated leaf samples using Trizol (Invitrogen, Carlsbad, California, USA), according to the manufacturer’s protocol. RNA quality and yield were determined by using Agilent 2100 Bioanalyzer (Agilent Technologies, Santa Clara, California, USA) and Qubit 2.0 Fluorometer (Invitrogen). Only RNA having integrity number (RIN) ≥ 8.0 were used. The cDNA libraries were multiplexed and then subjected to high throughput RNA-Seq using Illumina HiScanSQ (Illumina Inc., San Diego, California, USA). The sequenced samples (eight genotypes, six libraries each: I and NI for each time) produced forty-eight raw RNA-Seq libraries, corresponding to more than 50 million of 50 bp single-end reads per library.

### Transcriptome analysis

The raw data of each library were filtered using the software NGS-QC Toolkit [15] to remove bad quality, adaptors and contaminated sequences. The libraries were then aligned to the *Citrus sinensis* reference genome [16] using the Tophat2 (v2.1.14) [17] plus Cufflinks (v2.2.1)[18] approach. The identification of novel transcripts and isoforms was performed based on Cufflinks (v2.2.1) approach [18], using the reference transcriptome annotation of *Citrus sinensis* reference genome [16]. All identified genes, including annotated and novel isoforms, are available at ‘Download’ section. A total of 49,564 individual transcripts with an average transcript length of 1.5 Kbp, achieved a BUSCO completeness of ~95% with [S: 77.9%, D: 17.0%], F: 4.7%, M:0.4%, n: 430, according to the viridiplantae OrthoDB release 10 [19], and thus validating the CitrusKB as a comprehensive citrus transcriptome resource. These data were used in the gene expression analysis. In contrast, the *C. sinensis* and *C. clementina* transcriptomes available at Phytozome database [20] achieved a similar BUSCO completeness values, i.e. C: 94.2% [S: 52.1%, D: 42.1%], F: 4.9%, M: 0.9%, n: 430, and C: 98.6% [S: 72.6%, D: 26.0%], F: 1.2%, M: 0.2%, n: 430, respectively.

### Transcriptome annotation

The assembled transcripts and multi-species citrus transcriptome were annotated with the tools InterProScan (v5.29.68.0) [21], EggNOG (v4.5.1) [22] and Blast2GO (v5.0) [23]. The WEGO tool [24] was used to plot the GO annotations. Both the WEGO *GO Annotation file format (GAF)* file and EggNOG annotation results and further analysis are available for download in the CitrusKB ‘Download’ section. An overview of the Citrus transcriptome Distribution of euKaryotic Orthologous Group (KOG or NOGs) and Gene Ontology annotation is also available for visualization (Table 1, Figure 2).

**Table 1.** *Citrus sinensis* transcriptome: assembly and annotation status. Annotation type indicates the analyzed characteristic of the transcriptome, Amount indicates a size or a number of molecules.

| **Annotation type** | **Amount** |
| --- | --- |
| Total number of transcripts | 49,564 |
| Total size of transcripts | 73,991,731bp |
| GC% | 41.06 |
| Longest transcript | 16,810bp |
| Transcripts > 1kb | 29,599 |
| Mean transcript size | 1,493bp |
| Median transcript size | 1,241bp |
| Blast2GO Annotated Transcripts | 33,086 |
| GO Terms | 56,655 |
| -Biological Process | 18,271 |
| -Cellular Component | 12,090 |
| -Molecular Function | 26,294 |
| EggNOG Annotated Transcripts | 35,330 |
| -Unique GO terms | 6,265 |
| -Unique KEEG KOs | 3,541 |
| -Unique BiGG reactions | 284 |

**Figure 2.**
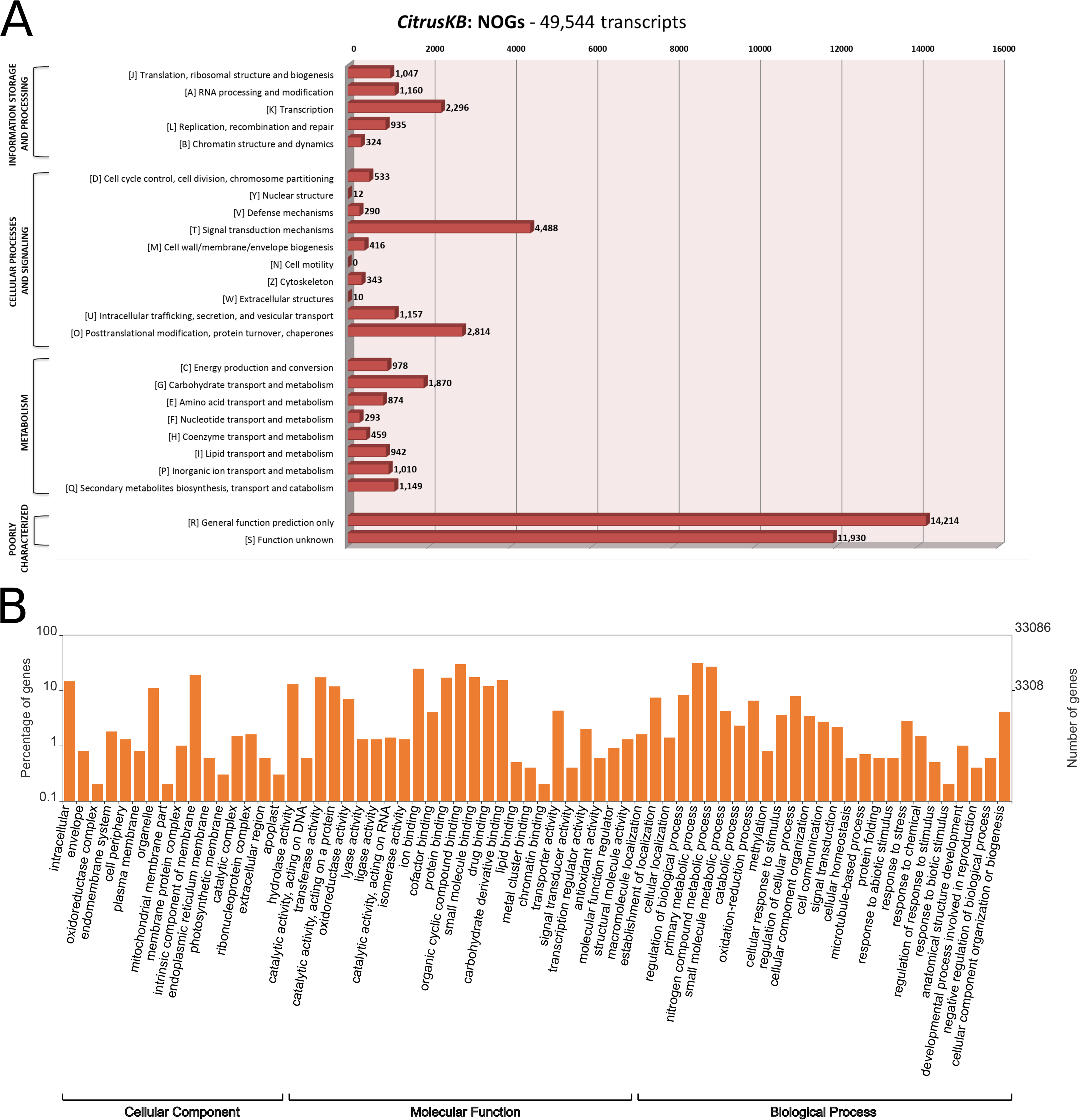
Annotation of *Citrus sinensis* CitrusKB transcriptome. EggNOG (A) and representative GO terms (B). The Y axis of GO annotations is presented in log(10) scale. Both, EggNOG and GO terms are fully searchable trough the Gene expression tables.

### Gene expression analysis

For differential gene expression analysis (DEG), the high quality reads used for transcriptome annotation step were mapped against the citrus reference genome using the STAR software (v2.4.1) [25]. The absolute number of aligned reads was extracted for each individual transcript from the created transcriptome which *loci* was associated to the reference genome, using a read counting approach over the mapped libraries. Further, the normalized expression variation with statistical significance analysis for each individual transcript was calculated between samples. The FDR (False Discovery Rate) correction [26] over the statistical significance found between the samples was applied to control false-positive significance transcript expression variation. These analysis were performed using DESeq, a Bioconductor R package [27]. The transcripts were considered as differentially expressed when statistical significance p-value was less than 0.05.

### Simple Sequence Repeats (SSR)

The search for Simple Sequence Repeats was performed using the tool SSRHTS software (version v1.0, unpublished). This approach performs custom SSR search using a reference sequence combined with high-throughput sequencing alignment data. This data is then used to validate detected SSRs and providing normalized occurrence values in a RPKM-based calculation for each SSR marker. Higher RPKM-based values denote higher expressed markers. The detected markers through all citrus genotypes, stored in a PostgreSQL database, describes the following information: sample name for each SSR occurred, sequence ID, sequence motif, start and end coordinates for SSR in reference sequence, motif length, length of repeat and RPKM-based value.

### Single Nucleotide Polymorphism (SNP)

To detect SNPs, RNA-Seq data were aligned to *Citrus sinensis* reference genome using the STAR aligner [25] with default parameters. Next, the genetic variants having expressed all alternate alleles were extracted using the SAMtools package [28]. The data generated were stored in the database and are represented by the following attributes: sequence ID, position, reference allele and SNP.

## Utility and Discussion

### CitrusKB implementation

CitrusKB was developed on a web-based environment and runs on a Linux operating system (Debian v7.7), which includes several common software packages: Apache HTTP server, PostgreSQL database, PHP, Java and Perl. The website (Figure 3) was developed using PHP, Java and HTML languages, and Twitter Bootstrap was used to achieve an enhanced user interaction. A Basic Local Alignment Search Tool (BLAST) [29] server is available in CitrusKB environment. As genome browser for transcriptome visualization, JBrowse [30] was implemented, showing all details of the transcripts and their alignments at nucleotide level.

**Figure 3.**
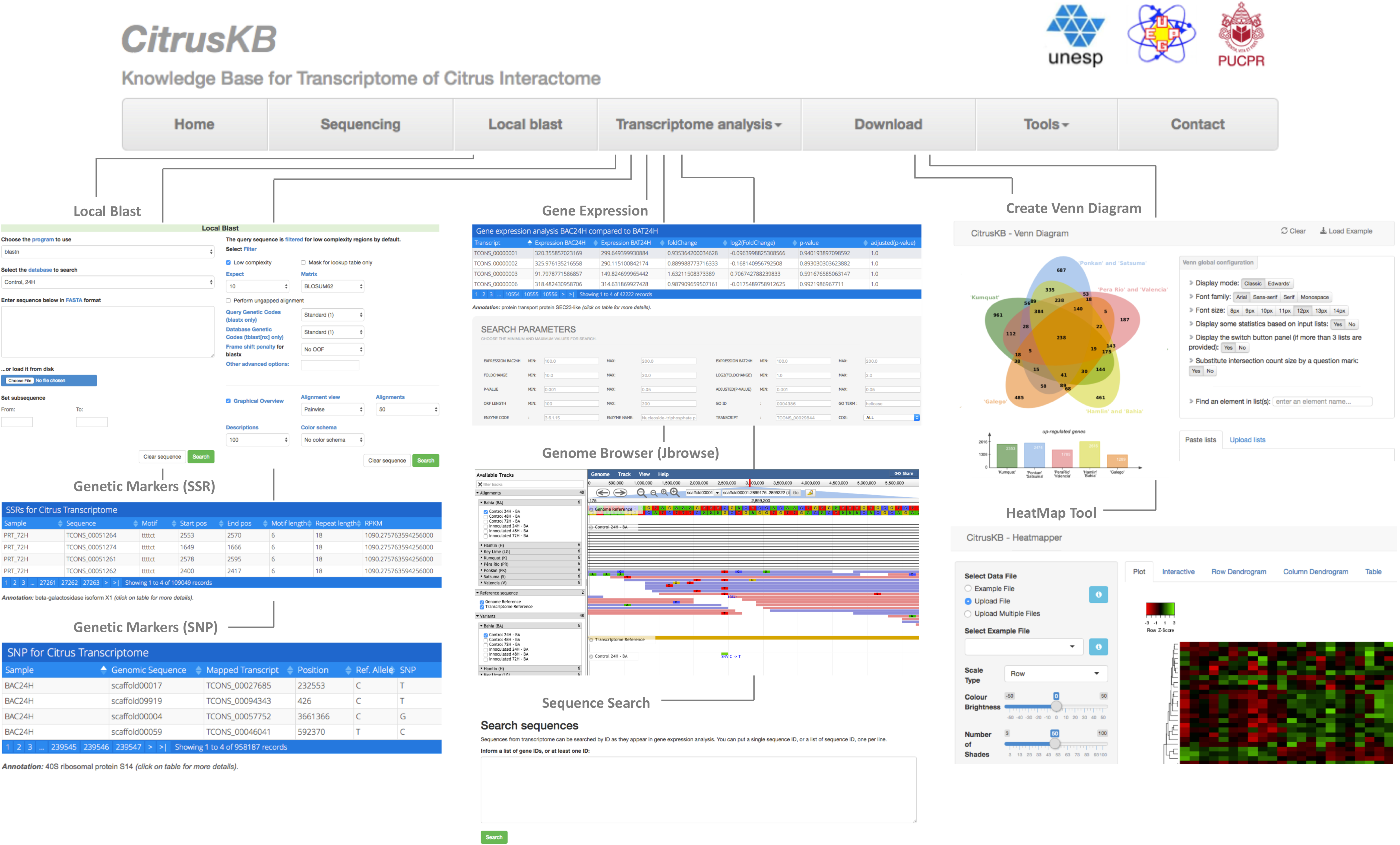
Citrus basic knowledge database (*CitrusKB*) website resources. Main functionalities are represented by the following screenshots: Gene Expression, Genome Browser, Sequence Search, Genetic Markers (SSR), Genetic Variants (SNP), Local Blast and transcriptome analysis tools Jvenn and Heatmapper.

### Local Blast

To provide an intuitive web-based graphical user interface (GUI) and a rapidly search in a large volume of data available in the CitrusKB, we used the standalone BLAST server (v2.2.26) [31]. To create the BLAST alignment database, we used the makeblastdb program of the NCBI BLAST (v2.2.30) software package [32]. Using web forms, users can perform nucleotide searches using the BLASTN tool and protein searches, using BLASTP tool.

### Sequence search

To perform searches for sequences from transcriptome, users can use as parameter a single or multiple sequence IDs, annotation keywords or gene names. The transcript sequence is retrieved by in-house, a script that performs the search over the stored transcriptome.

### Gene expression

In this section, users can view the gene expression profile of the sequenced libraries. For each citrus cultivar, differential analysis was performed to determine expression variation between pairs of libraries, including the following information: sequence ID, annotation, InterProScan [21] domains prediction, FPKM (Fragments per Kilobase of Exon per Million Fragments Mapped) per sample expression, Log_2_(Fold-change), p-value and FDR-corrected p-value. They are shown in a table that allows users to dynamically sort them. In addition, users can perform custom searches based on attributes of the analysis and limit the search by applying specific minimum and maximum filtering criteria for: gene expression values, fold-change alteration, p-value and FDR-corrected p-value. Filtered values can be exported to tables and users can download the results for additional analysis.

### Genome browser

CitrusKB uses JBrowse to dynamically browse and let users visualize, as multiple configurable tracks, the data corresponding to genome sequence, reference transcriptome, alignments information (sequencing coverage, splice junctions) and genetic variants (SNP, Insertions, Deletions). Available tracks are classified into three distinct categories, named as ‘Alignments’, ‘Reference Sequence’ and ‘Variants’. At ‘Reference Sequence’ category, users can visualize a reference genome track and a reference transcriptome track that provides a visualization of annotated information and the gene exon-intron structure. Users can also examine the expression level, as well as sequencing depth and coverage of each gene at the ‘Alignments’ category, including details related to alignment such as splice junctions and mismatched positions. All the forty-eight tracks were created from BAM alignment files to display short RNA-Seq reads aligned to the citrus reference genome. Users can also visualize single nucleotide variants for all tracks into ‘Variants’ category. The information is showed by JBrowse from VCF files.

### Genetic molecular markers

The SSR and SNP can be visualized by two distinct tables. In the first table, detected SSR in all samples are available for search. This table consists of sample ID, sequence ID, sequence, annotation, motif, start and end coordinates for SSR, motif and repeat length, and RPKM value. In the second table, SNPs are listed with the following attributes: sample ID, sequence name (scaffold from reference *C. sinensis* genome), position, reference allele and SNP. Both tables allow to dynamically sort the information and the user also can configure the search by choosing a specific citrus sample.

### Tools page

To facilitate the interpretation and to visualize the transcriptome annotation and differential expression analysis results, CitrusKB provides two different third-part commonly used tools for transcriptome analysis: (a) Venn diagrams, based on jvenn [33]; and (b) HeatMaps, implemented by the Heatmapper software [34]. Further details regarding on how to use these tools are shown in the topic “Using CitrusKB: a study case”.

### Download page

CitrusKB provides a user-friendly interface for the download of entire data sets. The HTTP links provide the download of all RNA-Seq raw sequencing reads from the forty-eight libraries, available in FASTA format and compressed as Gziped files. Users can also download the following data: reference genome, assembled reference transcriptome sequences and gene structure in GFF format, all the gene expression analysis, and other details of SSRs and SNPs genetic markers. The raw data is also available under BioProject number PRJNA470961.

## Using CitrusKB: a study case

The following section summarizes one example of CitrusKB usability. It is based on a study of differentially expressed genes (DEGs), revealing important transcripts found in the spatial and temporal citrus and *X. citri* interactome. The thresholds used during transcript search in the CitrusKB gene expression tables to consider a DEG were set as: fold-change ≥ 2 for up-regulated gene, fold-change ≤ −0.5 for down-regulated gene, both with an adjusted p-value ≤ 0.05. The resulting tables were exported in the “TSV” file format (‘export data’ button), and directly imported in the CitrusKB Venn Diagram Tool (‘Tools’ tab). The GAF file available in the ‘download section’ of the CitrusKB was used for the exploration of the enriched GO terms, with the use of the WEGO tool. In addition, the TSV file can be also imported in the CitrusKB HeatMap Tool (‘Tools’ tab) and interactively inspected in the form of heat maps. The annotation products, Gene Ontology terms and IDs, predicted NOGs and enzyme code of each DEG transcript can be retrieved through the gene expression tables or exported in the TSV file that can be further explored for other biological interpretations and/or downstream analysis.

It is worth to mention that the following study consist in a general example of the application of the CitrusKB which was not intended to show a comprehensive transcriptome analysis, but to illustrate the database use, the built-in tools to facilitate the transcriptome data integration, visualization and comparison. The objective to include this example was to present the power of the CitrusKB to revel the plant-pathogen related gene targets for further and more advanced studies. Researchers working with different plant-pathogen interaction topics can raise specific questions using the CitrusKB gene expression tables search engine and genome browser.

## A case of study: Global Overview of the DEG of the Citrus and *X. citri* interactome with identification of transcripts potentially related to plant-pathogen interactions

The analysis of the predicted DEGs of eight citrus genotypes in three-time stages (24 h, 48 h, and 72 h) revealed a plethora of DEG transcripts which can be strictly related with plant pathogen interaction, such as defense mechanisms and resistance and susceptibility to *X. citri*. In this analysis, the eight genotypes were grouped in five clusters: the first contains the highly resistant ‘Kumquat’ genotype; the second contains the resistant Mandarins ‘Ponkan’ and ‘Satsuma’; the third with the less resistant sweet oranges ‘Pera’ and ‘Valencia’; the fourth with the more susceptible genotypes ‘Hamlin’ and ‘Bahia’, and the fifth with the highly susceptible ‘Mexican’ lime genotype. In summary, ‘Kumquat’, ‘Ponkan and Satsuma’, ‘Hamlin’ and ‘Bahia’ stand as the genotypes with the larger number of up-regulated genes (average of 2,350), whereas ‘Kumquat’ exhibits the largest number of down-regulated genes (1,602) (Figure 4). The ‘Kumquat’ genotype also exhibits the largest number of exclusive up-regulated (961) and down-regulated genes (1,602). The list of the transcripts used to generate the Venn diagrams is provided as supplementary material (Table S1) for further exploitation.

**Figure 4.**
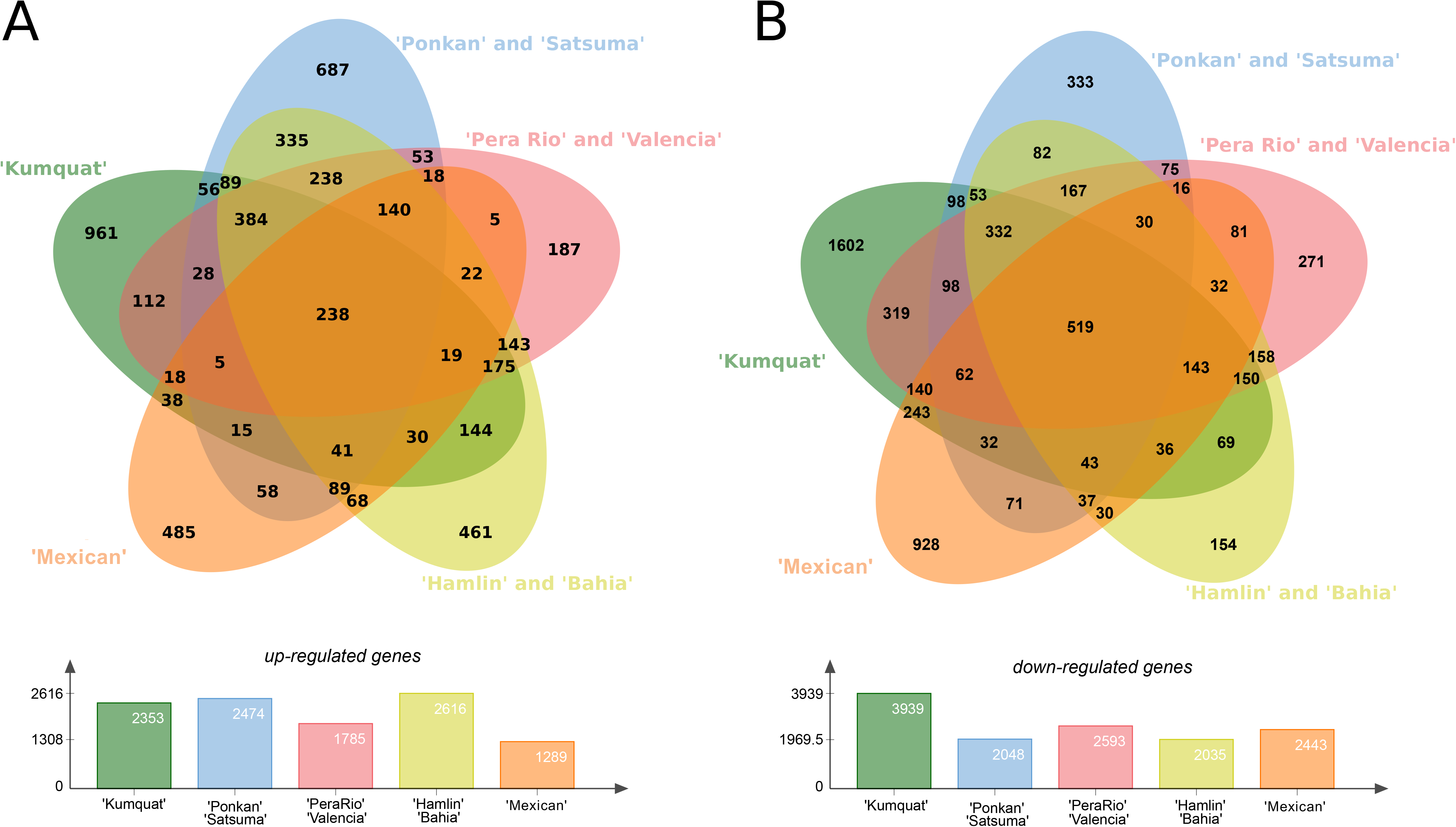
Differentially *expressed genes* (*DEGs*) present in the eight citrus genotypes considering the three analyzed time stages (24 h, 48 h and 72 h) pos *X. citri* inoculation. (A) Upper panel, Venn diagram with the Up-regulated genes; Lower panel, total number of DEGs for each analyzed genotype group. (B) Upper panel, venn diagram with the Down-regulated genes; Lower panel, total number of DEGs for each analyzed genotype group.

In general, the presented analysis also revealed 3,827 (7%) up-regulated and 4,889 (9%) down-regulated unique DEGs, and thus representing an overall pattern of *C. sinensis* response to *X. citri* infection. Among those, 642 up-regulated and 782 down-regulated transcripts correspond to hypothetical or uncharacterized proteins, and thus, may be considered as potential new targets for functional studies of plant-pathogen interaction mechanisms (Table S2). In addition, 2,307 up-regulated and 2,743 down-regulated transcripts are classified in NOGs (Table 2). The most prevalent DEGs NOGs belongs to ‘transcription’, ‘signal transduction mechanisms’, ‘posttranslational modification, protein turnover, chaperones’, ‘carbohydrate transport and metabolism’, and ‘secondary metabolites biosynthesis, transport and catabolism’. There are more than 40% of up-regulated DEGs related to ‘replication, recombination and repair’, and ‘cytoskeleton’ NOGs categories in comparison to down-regulated DEGs. In contrast, much more enriched down-regulated DEG were identified, such as those related to ‘transcription’, ‘cell wall/membrane/envelope biogenesis’, ‘carbohydrate transport and metabolism’, ‘coenzyme transport and metabolism’, ‘inorganic ion transport and metabolism’, and ‘secondary metabolites biosynthesis, transport and catabolism’.

**Table 2.** EggNOG classification of up- and down-regulated DEGs identified among the eight citrus genotypes in the all-time stages. Transcripts with only one NOG were considered (Table S1 for a comprehensive list of the transcripts and annotation).

| **EggNOG category** |  | **Up-regulated transcripts** | **Down-regulated transcripts** |
| --- | --- | --- | --- |
| **Information Storage and Processing** | [J] Translation, ribosomal structure and biogenesis | 91 | 96 |
|  | [A] RNA processing and modification | 28 | 53 |
|  | [K] Transcription | 179 | 261 |
|  | [L] Replication, recombination and repair | 72 | 22 |
|  | [B] Chromatin structure and dynamics | 7 | 7 |
| **Cellular Processes and Signaling** | [D] Cell cycle control, cell division, chromosome partitioning | 42 | 26 |
|  | [V] Defense mechanisms | 36 | 33 |
|  | [T] Signal transduction mechanisms | 552 | 566 |
|  | [M] Cell wall/membrane/envelope biogenesis | 24 | 69 |
|  | [Z] Cytoskeleton | 54 | 27 |
|  | [W] Extracellular structures | 2 | 2 |
|  | [U] Intracellular trafficking, secretion, and vesicular transport | 59 | 55 |
|  | [O] Posttranslational modification, protein turnover, chaperones | 271 | 280 |
| **Metabolism** | [C] Energy production and conversion | 92 | 112 |
|  | [G] Carbohydrate transport and metabolism | 184 | 283 |
|  | [E] Amino acid transport and metabolism | 101 | 99 |
|  | [F] Nucleotide transport and metabolism | 30 | 26 |
|  | [H] Coenzyme transport and metabolism | 23 | 88 |
|  | [I] Lipid transport and metabolism | 87 | 115 |
|  | [P] Inorganic ion transport and metabolism | 119 | 170 |
|  | [Q] Secondary metabolites biosynthesis, transport and catabolism | 141 | 245 |

These findings may suggest for a global set of genes potentially related to the plant response against the pathogen infection. For instance, among the identified NOGs (Figure 2A-2B), 440 out of 2296 (19%), 1118 out of 4488 (24%) and 551 out of 2814 (19%) represents to DEGs related to ‘transcription’, ‘signal transduction mechanisms’, and ‘posttranslational modification, protein turnover, chaperones’, respectively, which may regulate the citrus metabolism, through the inactivation and activation of genes in a putative attempt to control the infection caused by the pathogen. Conversely, 69 out of 290 (23%) DEGs related to ‘defense mechanisms’ were also identified, revealing several transcripts with detoxification related functions (Figure 2A, Table S2) which may be involved it the scavenge of the reactive oxygen species (ROS) generated by aerobic metabolism, and also by other processes, such as growth development, programmed cell death and response to biotic and abiotic environmental stimuli. All these processes can be linked to *X. citri* infection. Overall, these results summarized the use of CitrusKB to identify several transcripts (Table S2) with potential relationship to plant-pathogen interactions.

## Conclusions

The basic knowledge for transcriptome of Citrus interactome, CitrusKB, was created to provide and facilitate researches on citrus biology, in particular to the genes involved on plant-pathogen interaction. The CitrusKB also provides tools on a user-friendly WEB interface that allows researchers to visualize, search, analyze, and browse information regarding citrus and their interaction with the citrus canker pathogen *X. citri*. To the best of our knowledge, this is the first *in vivo* database for analysis of citrus infected by a pathogen, and we expect that it will bring a substantial contribution for researches in the area, particularly in the field of molecular plant-pathogen interactions. We will continue updating CitrusKB by incorporating novel literature data as they get available over time.

## Ethics approval and consent to participate

Not applicable.

## Consent for publication

Not applicable.

## Availability of data and material

The datasets generated and/or analyzed during the current study are available at the institutional website: http://bioinfo.deinfo.uepg.br/citrus.

## Authors’ contributions

AF, JBJ, HF, MITF, RPLJ, AMV, RHH, JAF designed the study (data generation and WEB-site); MMM, TDCGC, JSC, GP, JB prepared and generated the sequencing data; AF, MITF, HAP, AMV, RHH, JAF processed, analyzed and annotated the data; AF, HAP, AMV, RHH, JB, JAF wrote and revised the manuscript.

## Acknowledgements

We thank Applied Bioinformatics Laboratory (LBA) from Brazilian Agricultural Research Corporation (Embrapa/CNPTIA, Campinas, SP, Brazil) by providing computational and bioinformatics software support for data analysis.

## Supplementary materials

**Table S1. List of the transcripts from the CitrusKB used for the study case.** DEG are up-regulated (Table S2A) or down-regulated (Table S2B) in inoculated plants genotypes compared to relative controls. The abbreviation used within tables are as follow: KUMQUAT (Kumquat), PK (Ponkan), STS (Satsuma), VAL (Valencia), PR (Pera), HAM (Hamlin), BA (Bahia), MEXICAN (Mexican Lime).

**Table S2. Differentially expressed genes (DEG) classified as potential targets for functional studies of plant-pathogen interaction mechanism.** DEG are organized as: annotated up-regulated (Table S2A) or down-regulated (Table S2B) hypothetical proteins, uncharacterized up-regulated (Table S2C) or down-regulated (Table S2D) proteins, transcripts classified in euKaryotic Orthologous Group (NOG) that are up-regulated (Table S2E) or down-regulated (Table S2F).

## References

1. Li W, Song Q, Brlansky RH, Hartung JS. Genetic diversity of citrus bacterial canker pathogens preserved in herbarium specimens. Proc Natl Acad Sci U S A. 2007;104:18427–32.

2. Talon M, Gmitter FG. Citrus genomics. Int J Plant Genomics. 2008;2008:528361.

3. Civerolo EL. Bacterial canker disease of citrus. J Rio Gd Val Hortic Soc. 1984;37:127–46.

4. Rossetti VV. Manual Ilustrado de Doenças dos Citros. Piracicaba: Fealq/ Fundecitrus; 2001.

5. Behlau F, Amorim L, Belasque J, Bergamin Filho A, Leite RP, Graham JH, et al. Annual and polyetic progression of citrus canker on trees protected with copper sprays. Plant Pathol. 2010;59:1031–6.

6. Barbosa, J.C., Gimenes-Fernandes, N., Massari, C.A., Ayres A. Incidência e distribuição de cancro cítrico em pomares comerciais Estado de São Paulo e sul do Triângulo Mineiro. Summa Phytopathol. 2001;27:30–5.

7. de Carvalho SA, de Carvalho Nunes WM, Belasque J, Machado MA, Croce-Filho J, Bock CH, et al. Comparison of Resistance to Asiatic Citrus Canker Among Different Genotypes of *Citrus* in a Long-Term Canker-Resistance Field Screening Experiment in Brazil. Plant Dis. 2015;99:207–18. doi:10.1094/PDIS-04-14-0384-RE.

8. Khalaf A, Moore GA, Jones JB, Gmitter FG. New insights into the resistance of Nagami kumquat to canker disease. Physiol Mol Plant Pathol. 2007;71:240–50. doi:10.1016/J.PMPP.2008.03.001.

9. Cernadas RA, Camillo LR, Benedetti CE. Transcriptional analysis of the sweet orange interaction with the citrus canker pathogens Xanthomonas axonopodis pv. citri and Xanthomonas axonopodis pv. aurantifolii. Mol Plant Pathol. 2008;9:609–31.

10. Fu XZ, Gong XQ, Zhang YX, Wang Y, Liu JH. Different transcriptional response to Xanthomonas citri subsp. citri between kumquat and sweet orange with contrasting canker tolerance. PLoS One. 2012;7.

11. Terol J, Tadeo F, Ventimilla D, Talon M. An RNA-Seq-based reference transcriptome for Citrus. Plant Biotechnol J. 2016;14:938–50. doi:10.1111/pbi.12447.

12. Wang J, Chen D, Lei Y, Chang JW, Hao BH, Xing F, et al. Citrus sinensis Annotation Project (CAP): A comprehensive database for sweet orange genome. PLoS One. 2014;9.

13. Murata MM, Omar AA, Mou Z, Chase CD, Grosser JW, Graham JH. Novel plastid-nuclear genome combinations enhance resistance to citrus canker in cybrid grapefruit. Front Plant Sci. 2019;9:1858. doi:10.3389/fpls.2018.01858.

14. da Silva ACR, Ferro JA, Reinach FC, Farah CS, Furlan LR, Quaggio RB, et al. Comparison of the genomes of two Xanthomonas pathogens with differing host specificities. Nature. 2002;417:459–63. doi:10.1038/417459a.

15. Patel RK, Jain M. NGS QC Toolkit: a toolkit for quality control of next generation sequencing data. PLoS One. 2012;7:e30619. doi:10.1371/journal.pone.0030619.

16. Wu GA, Prochnik S, Jenkins J, Salse J, Hellsten U, Murat F, et al. Sequencing of diverse mandarin, pummelo and orange genomes reveals complex history of admixture during citrus domestication. Nat Biotechnol. 2014;32:656–62.

17. Kim D, Pertea G, Trapnell C, Pimentel H, Kelley R, Salzberg SL. TopHat2: accurate alignment of transcriptomes in the presence of insertions, deletions and gene fusions. Genome Biol. 2013;14:R36. doi:10.1186/gb-2013-14-4-r36.

18. Trapnell C, Williams BA, Pertea G, Mortazavi A, Kwan G, van Baren MJ, et al. Transcript assembly and quantification by RNA-Seq reveals unannotated transcripts and isoform switching during cell differentiation. Nat Biotechnol. 2010;28:511–5. doi:10.1038/nbt.1621.

19. Simão FA, Waterhouse RM, Ioannidis P, Kriventseva E V., Zdobnov EM. BUSCO: assessing genome assembly and annotation completeness with single-copy orthologs. Bioinformatics. 2015;31:3210–2. doi:10.1093/bioinformatics/btv351.

20. Goodstein DM, Shu S, Howson R, Neupane R, Hayes RD, Fazo J, et al. Phytozome: A comparative platform for green plant genomics. Nucleic Acids Res. 2012;40:D1178.

21. Quevillon E, Silventoinen V, Pillai S, Harte N, Mulder N, Apweiler R, et al. InterProScan: protein domains identifier. Nucleic Acids Res. 2005;33 Web Server issue:W116–20. doi:10.1093/nar/gki442.

22. Jensen LJ, Julien P, Kuhn M, von Mering C, Muller J, Doerks T, et al. eggNOG: automated construction and annotation of orthologous groups of genes. Nucleic Acids Res. 2007;36 Database:D250–4. doi:10.1093/nar/gkm796.

23. Conesa A, Götz S. Blast2GO: A Comprehensive Suite for Functional Analysis in Plant Genomics. Int J Plant Genomics. 2008;2008:1–12. doi:10.1155/2008/619832.

24. Ye J, Fang L, Zheng H, Zhang Y, Chen J, Zhang Z, et al. WEGO: A web tool for plotting GO annotations. Nucleic Acids Res. 2006;34 WEB. SERV. ISS.:W293.

25. Dobin A, Davis CA, Schlesinger F, Drenkow J, Zaleski C, Jha S, et al. STAR: ultrafast universal RNA-seq aligner. Bioinformatics. 2013;29:15–21. doi:10.1093/bioinformatics/bts635.

26. Benjamini Y, Hochberg Y. Controlling the False Discovery Rate: A Practical and Powerful Approach to Multiple Testing. J R Stat Soc Ser B. 1995;57:289–300.

27. Anders S, Huber W. Differential expression analysis for sequence count data. Genome Biol. 2010;11:R106. doi:10.1186/gb-2010-11-10-r106.

28. Li H, Handsaker B, Wysoker A, Fennell T, Ruan J, Homer N, et al. The Sequence Alignment/Map format and SAMtools. Bioinformatics. 2009;25:2078– doi:10.1093/bioinformatics/btp352.

29. Altschul SF, Gish W, Miller W, Myers EW, Lipman DJ. Basic local alignment search tool. J Mol Biol. 1990;215:403–10.

30. Buels R, Yao E, Diesh CM, Hayes RD, Munoz-Torres M, Helt G, et al. JBrowse: a dynamic web platform for genome visualization and analysis. Genome Biol. 2016;17:66.

31. Johnson M, Zaretskaya I, Raytselis Y, Merezhuk Y, McGinnis S, Madden TL. NCBI BLAST: a better web interface. Nucleic Acids Res. 2008;36 Web Server issue.

32. Camacho C, Coulouris G, Avagyan V, Ma N, Papadopoulos J, Bealer K, et al. BLAST+: architecture and applications. BMC Bioinformatics. 2009;10:421.

33. Bardou P, Mariette J, Escudié F, Djemiel C, Klopp C. jvenn: an interactive Venn diagram viewer. BMC Bioinformatics. 2014;15:293. doi:10.1186/1471-2105-15-293.

34. Babicki S, Arndt D, Marcu A, Liang Y, Grant JR, Maciejewski A, et al. Heatmapper: web-enabled heat mapping for all. Nucleic Acids Res. 2016;44:W147–53. doi:10.1093/nar/gkw419.

